# Expression and Neurotransmitter Association of the Synaptic Calcium Sensor Synaptotagmin in the Avian Auditory Brain Stem

**DOI:** 10.1101/2021.10.01.462838

**Authors:** Katrina M. MacLeod, Sangeeta Pandya

**Author notes:** Corresponding Author: Katrina MacLeod, Department of Biology, University of Maryland, College Park, MD 20742.

## Abstract

In the avian auditory brain stem, acoustic timing and intensity cues are processed in separate, parallel pathways via the two division of the cochlear nucleus, nucleus angularis (NA) and nucleus magnocellularis (NM). Differences in excitatory and inhibitory synaptic properties, such as release probability and short-term plasticity, contribute to differential processing of the auditory nerve inputs. We investigated the distribution of synaptotagmin, a putative calcium sensor for exocytosis, via immunohistochemistry and double immunofluorescence in the embryonic and hatchling chick brain stem (*Gallus gallus*). We found that the two major isoforms, synaptotagmin 1 (Syt1) and synaptotagmin 2 (Syt2), showed differential expression. In the NM, anti-Syt2 label was strong and resembled the endbulb terminals of the auditory nerve inputs, while anti-Syt1 label was weaker and more punctate. In NA, both isoforms were intensely expressed throughout the neuropil. A third isoform, synaptotagmin 7 (Syt7), was largely absent from the cochlear nuclei. In nucleus laminaris (NL, the target nucleus of NM), anti-Syt2 and anti-Syt7 strongly labeled the dendritic lamina. These patterns were established by embryonic day 18 and persisted to postnatal day 7. Double labeling immunofluorescence showed Syt1 and Syt2 were associated with Vesicular Glutamate Transporter 2 (VGluT2), but not Vesicular GABA Transporter (VGAT), suggesting these Syt isoforms were localized to excitatory, but not inhibitory, terminals. These results suggest that Syt2 is the major calcium binding protein underlying excitatory neurotransmission in the timing pathway comprising NM and NL, while Syt2 and Syt1 regulate excitatory transmission in the parallel intensity pathway via cochlear nucleus NA.

## Introduction

Synaptic connections between auditory nerve axons and their neuronal targets in the cochlear nuclei define how acoustic information is processed and transmitted along the ascending pathways. Differential regulation of these properties may depend on differences in the molecular components underlying presynaptic mechanisms of transmitter release.

Synaptotagmins are a family of transmembrane Ca^2+^-binding proteins whose two main isoforms, Syt1 and Syt2, are thought to be the major calcium sensors for neurotransmitter exocytosis (Südhof 2013; Wolfes and Dean 2020). Among 17 different family members, synaptotagmin 1 (Syt1) and synaptotagmin 2 (Syt2) have been shown to drive fast exocytosis at excitatory and inhibitory synapses in mammalian neocortex, hippocampus, cerebellum and brain stem (Geppert et al. 1994; Fernández-Chacón et al. 2001; Pang et al. 2006; Xu et al. 2007; Chen et al. 2015; Kochubey et al. 2016). Expression studies have shown that the isoforms are differentially expressed and in some cases are subregion or layer specific, suggesting synapse specific regulation of targeting of each subtype (Fox and Sanes 2007; Mittelsteadt et al. 2009). The Syt1 isoform appears predominant in the cortex and other forebrain, while the Syt2 isoform is the predominant form in the hindbrain, where isoform expression may also be developmentally regulated (Kochubey et al. 2016). Genetic knockout rescue studies suggest that Syt1 and Syt2 can drive fast exocytosis interchangeably (Stevens and Sullivan 2003; Nagy et al. 2006; Xu et al. 2007), but subtle differences in kinetics or Ca+2 binding affinities may partially explain the synapse specificity (Sugita et al. 2002; Hui et al. 2005; Chen et al. 2017a). Furthermore, different forms of exocytosis may rely on different isoforms, with synaptotagmin 7 (Syt7) being a key candidate involved in asynchronous release and short-term synaptic facilitation (Wen et al. 2010; Bacaj et al. 2013; Jackman et al. 2016; Chen et al. 2017b; Turecek and Regehr 2018; Deng et al. 2020).

The auditory brain stem is a useful model system for understanding how synaptic function relates to neural coding. In the avian and mammalian auditory brain stem, short-term depression and facilitation are differentially expressed in parallel pathways (MacLeod and Carr 2007; MacLeod et al. 2007; Cao and Oertel 2010). Auditory nerve axons bifurcate and form morphologically and physiologically distinct synapse types onto their targets in the cochlear nucleus, suggesting target-dependent regulation of synaptic properties (Jhaveri and Morest 1982; Carr and Boudreau 1993). The auditory nerve fibers (ANFs) form large, calyceal synapses in the avian cochlear nucleus magnocellularis (NM) that have tightly synchronized, highly reliable release properties and profound short-term synaptic depression (Trussell 1999). Smaller, bouton-like synapses from ANFs onto neurons in the sister cochlear nucleus angularis (NA) have small amplitude excitatory currents and mixed facilitation and depression short-term synaptic plasticity (MacLeod and Carr 2005; MacLeod et al. 2007). Finally, the avian cochlear nucleus (CN) receives descending feedback inhibitory synapses that show substantial asynchronous release (Monsivais et al. 2000a; Burger et al. 2005; Tang and Lu 2012). Whether these cell-type specific differences in synaptic release properties are determined by the specific molecular isoforms involved in exocytosis is unknown. To investigate the molecular basis of calcium triggered release, we measured the expression and distribution of Syt1, Syt2 and Syt7 proteins via immunohistochemistry and double immunofluorescence in the brain stems of embryonic and hatchling chick (*Gallus gallus*). Our results suggest Syt2 is the dominant isoform underlying excitatory neurotransmission in the avian auditory brain stem circuits. In contrast, the Syt1 and Syt7 isoforms are less prominent but may be differentially involved in establishing or modulating synaptic pathways in the avian auditory brain stem.

## Results

To investigate the expression patterns of the synaptotagmin isoforms in the avian auditory brain stem nuclei, we performed immunohistochemistry using antibodies against syt1, syt2 and syt7 in paraformaldehyde fixed tissue. All animals used for immunohistochemistry and immunofluorescence were chicks perfused within 24 hours of hatching, except for the developmental series which used chick embryos aged embryonic day (E)18, just hatched chicks (posthatch day (P) 0 or 1), or week-old (P7) chicks. We confirmed labeling for Syt1, Sty2 and Syt7 with expected patterns outside of the auditory brain stem nuclei: dense labeling in the medial vestibular nuclei (MVN) and known patterning of synaptotagmin isoforms in the cerebellum (Popratiloff et al. 2003; Fox and Sanes 2007; Shao et al. 2012).

### Synaptotagmin 2 was abundant throughout the avian auditory brainstem

An antibody against synaptotagmin 2 (znp-1, hereafter ‘anti-Syt2’; Table 1) revealed abundant expression throughout the auditory brain stem nuclei (Fig. 1A). Anti-Syt2 label was similarly abundant in both cochlear nuclei NM and NA, as well as the target nucleus for NM involved in interaural timing difference coding, nucleus laminaris (NL). Anti-Syt2 label showed distinct patterns of expression associated with known synaptic terminal fields within the auditory nuclei, resembling presynaptic terminals, but was weak or absent from the principal neuronal cell bodies or the axonal projections. For example, in NM the label formed large, continuous shapes that wrapped around the negative NM cell bodies and thus resembled the end bulb terminals of the auditory nerve fibers (ANF)(Fig. 1B). In contrast, the label in NA was dense but punctate, located in the neuropil but largely absent from apparent cell bodies (Fig. 1C). In medial NL, the labeled puncta in the dorsal and ventral dendritic fields were so dense as to be difficult to resolve (Fig. 1D), but in lateral NL individual puncta were distinguishable (Fig. 1E). The anti-Syt2 label in NL appeared equally abundant in dorsal and ventral dendritic lamina, with weaker label in the central cell body region in a similar pattern to synaptic vesicle protein 2 (Parameshwaran et al. 2001; MacLeod et al. 2006).

**Table 1.** Primary Antibody List

| Antibody | Immunogen | Manufacturer and type |
| --- | --- | --- |
| Synaptotagmin-1 | Rat synaptic junction complex, cytoplasmic domain | DSHB mouse monoclonal mab-48 |
| Synaptotagmin-2 | Zebrafish embryo full length protein | DSHB mouse monoclonal znp-1 |
| Synaptotagmin-7 | Recombinant protein from rat SYT7 (AA 46 to 133) | Synaptic Systems Cat#105173 rabbit polyclonal |
| Vesicular glutamate transporter 2 | Recombinant protein from rat VGLUT2 (AA 510 to 582) | Synaptic Systems Cat#135403 rabbit polyclonal |
| Vesicular GABA transporter | Rat VGAT C-terminus cytoplasmic domain | Alpha Diagnostics VGAT11A rabbit polyclonal |

DSHB: Developmental Studies Hybridoma Bank

**Figure 1.**
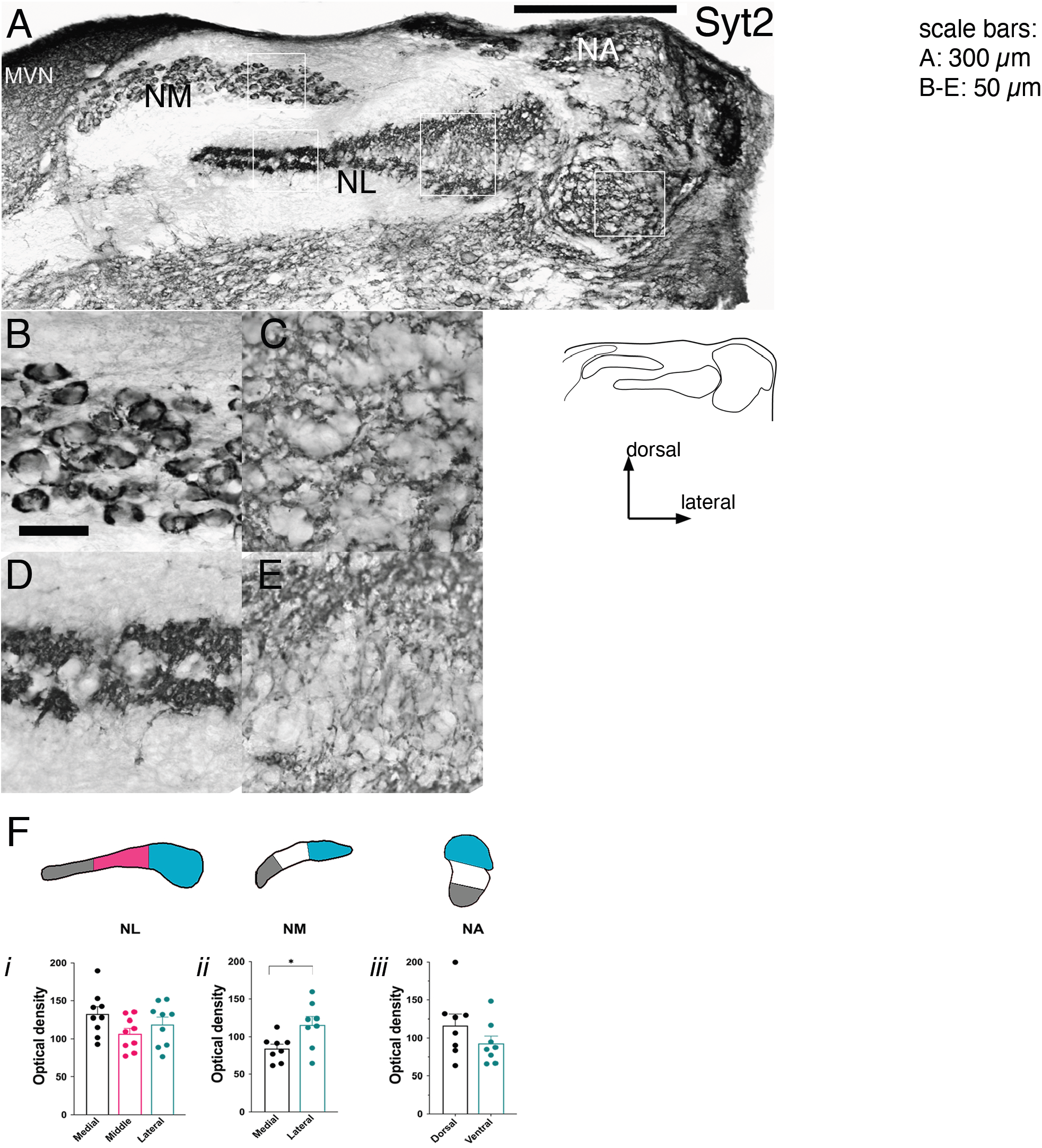
Anti-Syt2 (znp-1) label showed abundant expression in the avian auditory brainstem nuclei in hatchling (P1) chick. A) Low power montage of dorsal right quadrant of transverse brain stem section. Dense label observed in NM, NL and NA, as well as MVN, but absent from fiber tracts surrounding NM and NL. White boxes indicate location of panels B-E. Schematic below shows outlines of nuclei. Scale bar: 300 µm. B) Dense label surrounded NM cell bodies, resembling axosomatic endbulb terminals. Scale bar: 50 µm, applies to panels B-E. C) NA showed anti-Syt2 label throughout the neuropil, surrounding negative cell bodies. D) Dense anti-Syt2 label were found in the dorsal and ventral dendritic fields of the medial NL. E) Dispersed anti-Syt2 puncta were found in the dorsal and ventral dendritic fields of lateral NL. F) Optical density analysis of anti-Syt2 label across the tonotopic axes of NL (i), NM (ii) and NA (iii). Each nucleus was divided into three regions of interest and the average OD measured. Syt2 label was slightly more abundant in lateral (lower best frequency) NM than medial (higher best frequency) NM (N = 7 sections, P = 0.0391, Wilcoxon matched-pairs signed rank test). No significant tonotopic variation in Syt2 label was seen between regions in NL (mediolateral dimension) or NA (dorsoventral dimension). Pooled data across 4 subjects.

### Synaptotagmin 1 was sparsely expressed in timing nuclei but abundant in cochlear nucleus angularis

Anti-synaptotagmin 1 (mab48, hereafter ‘anti-Syt1’, Table 1) immunohistochemistry revealed differential expression across the auditory brain stem nuclei (Fig. 2A). Anti-Syt1 label in NM was comparatively sparsely present in small puncta surrounding cell bodies (Fig. 2B) rather than continuous endbulb-like profiles like the anti-Syt2 label. In lateral NM some cell bodies showed a complete absence of anti-Syt1 label. In contrast, anti-Syt1 label was abundant in NA forming a dense web of darkly labeled puncta (Fig. 2C). In NL, anti-Syt1 label was even weaker than in NM, especially in the medial and central areas where only a few sparse puncta were observed (Fig. 2D), although Syt1-positive puncta were more abundant in the dendritic fields of lateral NL (Fig. 2E). We were also able to observe heavy anti-Syt1 labeling within the boundaries of the superior olivary nucleus (SON)(Fig. 2F).

**Figure 2.**
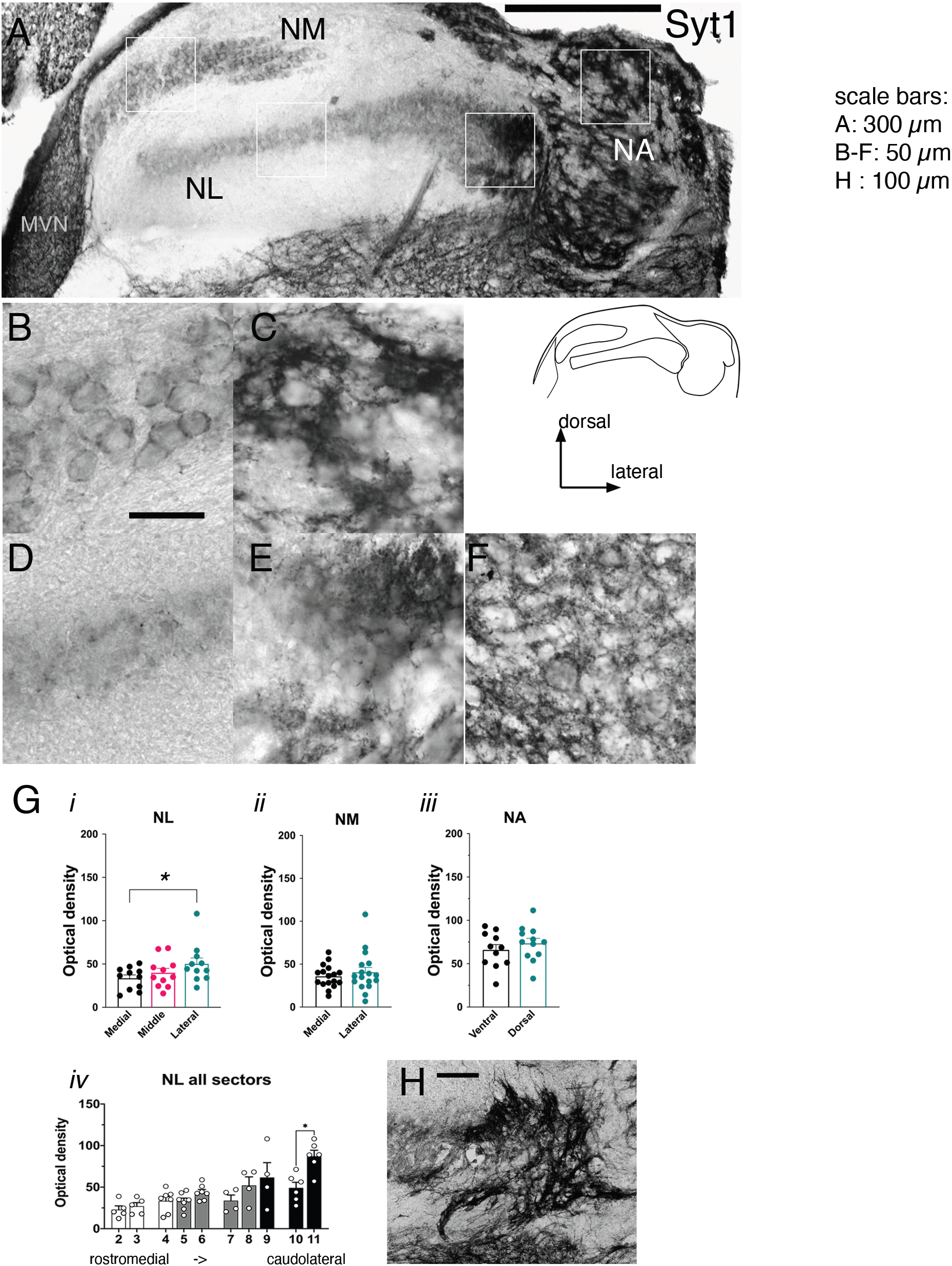
Anti-Syt1label was heterogeneous among the auditory nuclei in the hatchling (P1) chick. Panel organization similar to Figure 1. A) Low power montage of dorsal right quadrant of transverse brain stem section showed weak labeling the timing nuclei NM and NL, but abundant labeling in NA. White boxes indicate location of panels B-E. Schematic below shows outlines of nuclei. Scale bar: 300 µm. B) Anti-Syt1 label in NM was punctate surrounding cell bodies but did not resemble endbulbs. Scale bar: 50 µm, applies to panels B-F. C) Dense labeling of the neuropil throughout NA. D) Anti-Syt1 label in medial NL was found in sparse puncta. E) Lateral region of NL showed dense anti-Syt1 label. F) Syt1 label was also evident in the SON (different section). G) Optical density analysis of Syt1 label across the tonotopic axes. (*i*) Analysis of NL divided into mediolateral regions showed a significant, though small, tonotopic effect (1-way ANOVA, N=11 sections (4 subjects), F (2, 20) = 4.546, P=0.0236; medial vs lateral, P=0.0218, Holm-Šídák’s multiple comparison test). No tonotopic effects were observed in NM (*ii*) or NA (*iii*). (*iv*) A more detailed sector-level analysis of Syt1 OD showed a significant effect in the caudolateral extreme of NL (sector 10 vs 11, Wilcoxon matched-pairs ranked test, N = 6 sections (3 subjects), P = 0.0312). All statistical comparisons were made within section. H) Extreme caudolateral area of NL corresponding to sector 11 showed anti-Syt1 labeling of fiber innervation. Scale bar: 100 µm.

Unlike Syt2 labeling, which appeared restricted to axonal terminals in the auditory nuclei, Syt1 labeling could sometimes be observed in fibers projecting to nucleus laminaris (Fig. 2H). These were not originating from NM; in one case, an isolated individual neuron located dorsally to NL dendritic field could be observed that appeared to ramify NL (Fig. 3). Other projection fibers entered caudolateral NL as a large bundle from the caudolateral margin but could not be traced further due to the density of anti-Syt1 label in the ventral brain stem (Fig. 2H).

**Figure 3.**
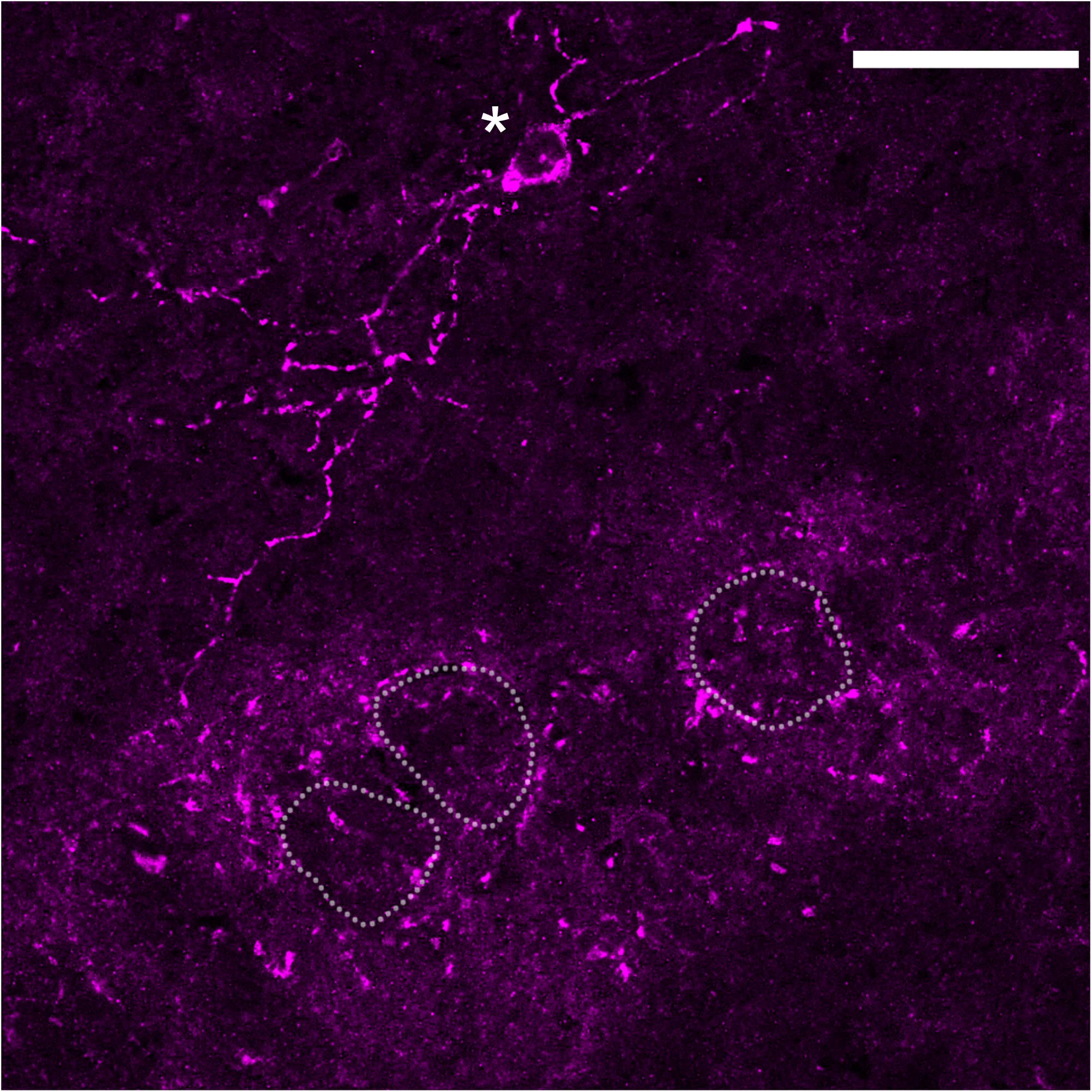
Anti-Syt1 labeling revealed fibers innervating NL from a dorsally located local neuron (asterisk). Dotted lines outline the cell body layer of NL. Scale bar: 30 µm.

### Synaptotagmin 7 is differentially expressed in auditory nuclei

Expression patterns observed using anti-synaptotagmin 7 (anti-Syt7) antibodies were distinct from both anti-Syt1 and anti-Syt2 labeling(Fig. 4A). Both cochlear nuclei NM and NA were weakly and diffusely labeled compared to NL, the MVN, or the ventral brain stem tissue (Fig. 4B-C). Like the other two isoforms, the anti-Syt7 label in NL appeared concentrated in the dendritic fields (Fig. 4D). The SON also showed abundant anti-Syt7 label (Fig. 4E).

**Figure 4.**
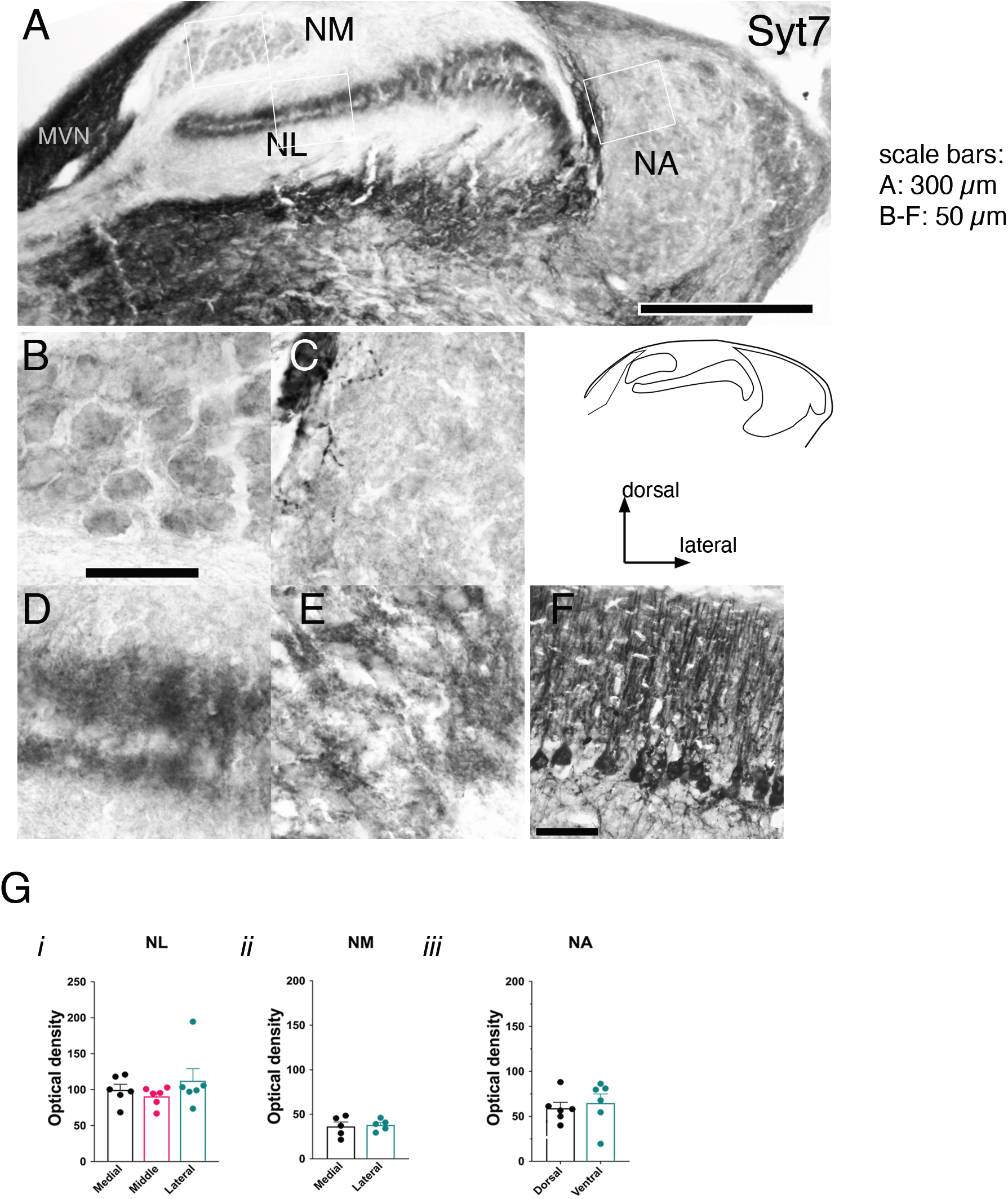
Anti-Syt7 label was heterogeneous among the auditory nuclei. A) Low power montage of dorsal right quadrant of transverse brain stem section showed weak labeling of the cochlear nuclei NA and NM, while there was robust labeling of NL dendritic fields, as well as MVN and non-auditory brain stem tissue. Panel organization similar to Figs. 1 and 2. White boxes indicate location of panels B-E. Schematic below shows outlines of nuclei. Scale bar: 300 µm. B) Anti-Syt7 label in NM was weak and punctate where present. Scale bar: 50 µm, applies to panels B-E. C) Anti-Syt7 was nearly absent from NA, although a few densely labeled fibers of unknown origin penetrated medial margins. D) In NL, diffuse labeling was seen in the dendritic lamina. E) Punctate anti-Syt7 label was abundant in the SON (different section). F) Anti-Syt7 robustly labeled Purkinje cell bodies and their apical dendrites in the cerebellum. Scale bar: 50 µm. G) Optical density analysis of Syt7 expression showed no variation across the tonotopic axes of NL (*i*), NM (*ii*) or NA (*iii*).

### Synaptotagmin expression patterns relative to the tonotopic axes

Avian auditory brain stem nuclei contain maps of best sound frequency (BF) and many cellular and molecular properties show systematic changes or gradients of expression along the tonotopic axis (see Discussion). To determine if the expression of synaptotagmins varied with the tonotopic axis, we analyzed the optical density of anti-Syt1, -Syt2 and -Syt7 label across the relevant orientation axes of the auditory nuclei. Optical density measurements were made in the medial versus lateral regions in NM and the dorsal versus ventral regions of NA (high versus low BF for each nucleus, respectively) (schematics shown in Fig. 1F). Optical density was also measured across the mediolateral length of NL, sampling in medial, middle and lateral regions, which correspond to high, middle and low BF regions. The cochlear nuclei showed no tonotopic expression gradients for any synaptotagmin, except for a small increase in density in the lateral region of NM for anti-Syt2 (Fig. 4F*ii*-*iii*, Wilcoxon matched pairs signed rank test, n = 7 pairs, P = 0.0391). For anti-Syt2 and anti-Syt1 label, expression in NL was more varied. Anti-Syt2 labeling was evenly distributed across the mediolateral dimension of NL with a slight dip in the middle region (Fig. 1F*i*; 1-way ANOVA P = 0.0476; multiple comparison test, adjusted P = 0.055, middle sector vs lateral sector). In contrast, for the anti-Syt1 label a large change in density was clearly observable across the mediolateral dimension of NL (Fig. 2G*i*; 1-way ANOVA P = 0.0236; post-hoc multiple comparison test showed a significant difference between medial vs lateral NL, P = 0.0218). The densest anti-Syt1 label labeling was observed in the extreme lateral area of NL (e.g. see Fig. 2A). To examine this pattern more accurately relative to the rostromedial to caudolateral direction of orientation of the BF mapping in NL (Rubel and Parks 1988), we further analyzed anti-Syt1 labeling by subsectors according to the rostromedial to caudolateral sector divisions (see Methods; c.f. Kuba et al. 2005). The mediolateral trends were observable in the middle sectors (4-6, 7-9; these pooled values are plotted in Fig. 2G*i*) but there was a significant increase in density in the most caudolateral sector (Fig 2H; sector 10 vs 11: Wilcoxon matched pairs signed rank text, n=6 pairs, P = 0.0312). Thus, Syt1 expression was preferentially observed in the NL sectors that corresponded to the lowest BF. In contrast, anti-Syt7 labeling was uniformly distributed across NL (Fig 4F*i*; 1-way ANOVA, P = 0.2189). These data suggest that the only isoform expressed with a notable tonotopic effect was Syt1, and then only in NL.

### Synaptotagmin expression patterns over development

Synapses in the chicken auditory cochlear nucleus are established and become functional prior to E17, then undergo morphological and physiological refinement up to hatching at E21 and several days thereafter (Jackson et al 1982; Brenowitz and Trussell 2001; see Kubke & Carr 2000 for review). To determine whether the synaptotagmin expression patterns observed in just-hatched chicks were present at earlier stages and persisted later, we repeated immunohistochemical experiments in parallel at three stages: late embryonic (“E18”), hatchling (“P1”) and week-old chicks (“P7”)(see Methods). Our data showed that Syt2, the dominant isoform, showed small increases in the density of label over this age range of development. Anti-Syt2 label, already dense at E18, increased with age (Fig. 5Ai, Bi; 2-way ANOVA main effect by age, F(2,113) = 5.109, P = 0.0075); there were significant differences between E18 and P1 (Tukey’s multiple comparison test, P = 0.00224) and between E18 and P7 (P = 0.0178). Comparing individual nuclei across ages, NL showed a significant increase between E18 and P7 (Fig. 5Bi, P = 0.0261).

**Figure 5.**
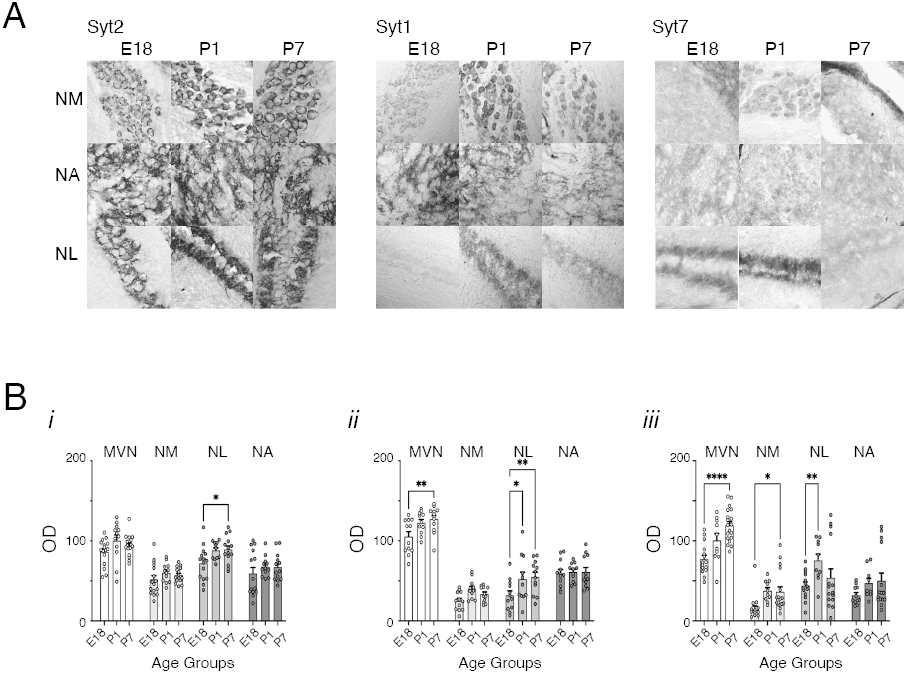
Synaptotagmin protein expression showed small but significant developmental changes across the auditory brain stem. A) Representative images showing developmental changes in expression of Syt2 (left panel), Syt1 (middle panel) and Syt7 (right panel) in NM (top row in each set of panels), NA (middle row) and NL (bottom row). Columns within each panel shows representative images per age group: E18, hatchlings (P1), or week-old chicks (P7). B) Summary of optical density changes with development for Syt2 (i), Syt1 (ii) and Syt7 (iii). Mean OD was measured from the whole nucleus as a region of interest in each section, pooled across 3 or 4 subjects for each nucleus and age (see Methods). Statistically significant effects of age were observed for anti-Syt2 (2-way ANOVA, main effect by age: F (6, 154) = 0.3953, P = 0.0012), anti-Syt1 (F (2, 128) = 9.365, P = 0.0002), and anti-Syt7 (F (2, 147) = 14.91, P < 0.0001). Asterisks indicate significant Tukey’s multiple comparisons tests (*, P < 0.05, **, P < 0.01, ****, P < 0.0001). Significant effects by nucleus were also found (see text).

Although brain stem regions are difficult to compare because they have different morphological structures and neuropil, we did observe differences in relative levels of Syt2 expression by nucleus (second main effect by nucleus, 2-way ANOVA, including MVN: F(3,154)=38.24, P < 0.0001; auditory only: F(2, 113) = 24.01, P < 0.0001). Expression in NL was significantly more dense compared to NM at all ages (at age E18: P = 0.0099; P1: P = 0.0015; P7: P < 0.0001; pairwise multiple comparisons Tukey’s test) and compared to NA after hatching (E18: P = 0.23; P1: P = 0.036; P7: P = 0.0086). There were no significant differences in Syt2 expression between NA and NM at any age. The cochlear nuclei had weaker expression of anti-Syt2 label at all ages than the MVN (MVN vs NM, all ages: P < 0.0001; MVN vs NA, at E18: P = 0.003; at P1 and P7: P < 0.0001) but NL and MVN had comparable expression.

Limited age-related changes in anti-Syt1 expression in the auditory nuclei were also observed (Fig. 4Aii, Bii; 2-way ANOVA main effect by age, F (2, 96) = 5.14, P = 0.0076); there were significant differences between E18 and P1 (multiple comparison Tukey’s test, P = 0.0014) and between E18 and P7 (P = 0.0245). Comparing individual nuclei across ages, NL showed a significant increase between E18 and P1 (P = 0.0215) and between E18 and P7 (P = 0.0053)(Fig. 4Bii). The MVN also showed an age-dependent increase between E18 and P7 (2-way ANOVA including MVN, main effect by age, F(2, 128) = 9.365, P = 0.0002, multiple comparison test for MVN E18 versus P7, P = 0.0084).

The auditory nuclei showed strikingly weaker expression of Syt1 relative to the MVN, as well as to the ventral brain stem areas overall (see Fig. 2). There was also a clear gradation in Syt1 label density among the auditory nuclei across ages, with NA>NL>NM (second main effect by nucleus, 2-way ANOVA, including MVN: F(3,128) = 158.6, P < 0.0001; auditory only: F(2, 96) = 21.79, P < 0.0001). In the late embryos, anti-Syt1 expression levels were significantly higher in NA relative to NM and NL (E18: NA vs NM, P < 0.0001, NA vs NL, P = 0.0013) while expression in NM and NL was comparable (P = 0.7433). By P7, NL anti-Syt1 label was not significantly less dense than NA and both NA and NL had significantly denser label than NM (P7, NA vs NL, P = 0.845, NM vs NL, P = 0.0162, NA vs NM, P < 0.0011).

Among the synaptotagmin isoforms, anti-Syt7 label showed weaker expression in the auditory nuclei but more robust developmental effects. Similar to anti-Syt1 and -Syt2 label, anti-Syt7 label showed a significant main effect by age (Fig. 4Aiii, Biii; 2-way ANOVA main effect by age, F (2, 106) = 7.341, P = 0.0010); there were significant differences between E18 and P1 (P = 0.0015) and between E18 and P7 (P = 0.0114). Anti-Syt7 label showed a significant effect of age in NM (E18 vs P7: P = 0.038) as well as in NL (E18 vs P1: P = 0.0092). In contrast, the anti-Syt7 label in NA showed no changes with development. The largest increases in anti-Syt7 label were seen in MVN (E18 vs P7: P < 0.001).

There were differences in the density of anti-Syt7 label between the auditory nuclei, with the sparsest label in NM whose OD measurements in embryonic tissue were often close to background. In comparison, Syt7 was significantly more densely expressed in NL (2-way ANOVA main effect by nucleus, F (3, 147) = 56.21, P < 0.001; at age E18: P = 0.0105; P1: P = 0.0059; Tukey’s multiple comparison tests). Density values for NA fell between NM and NL, with no significant differences. Anti-Syt7 expression in MVN was more dense than in all three auditory nuclei, increasingly so with age (at age P7, MVN > NA, NM and NL, P < 0.001). In summary, all three synaptotagmin isoforms show small developmentally related changes in protein expression that nevertheless did not substantially change the patterns of expression among the auditory nuclei.

### Synaptotagmin 2 was associated with excitatory synaptic terminals

The anti-Syt2 label strongly resembled the endbulbs in NM, which are the large axosomatic terminals of auditory nerve axons that provide strong excitatory input to the NM principal cells. To determine whether the synaptotagmin label was associated with excitatory presynaptic terminals, we performed double immunofluorescent histochemistry with antibodies against Syt1 (or Syt2) and against vesicular glutamate transporter (VGluT2), a protein responsible for transmitter loading of synaptic vesicles in glutamatergic terminals. In NM, both anti-Syt2 and anti-VGluT2 label resembled end bulb terminals (Fig. 6A, A’) and nearly completely co-localized (Fig. 6A”). In NA (Fig. 6C) and NL (Fig. 6E), labeling was punctate for both antibodies and also showed nearly completely overlap in the neuropil of NA and dendritic field of NL. In the cell body layer of NL, however, anti-VGluT2 was largely excluded, and some Syt2+/VGluT2-profiles could be observed (Fig. 6e).

**Figure 6.**
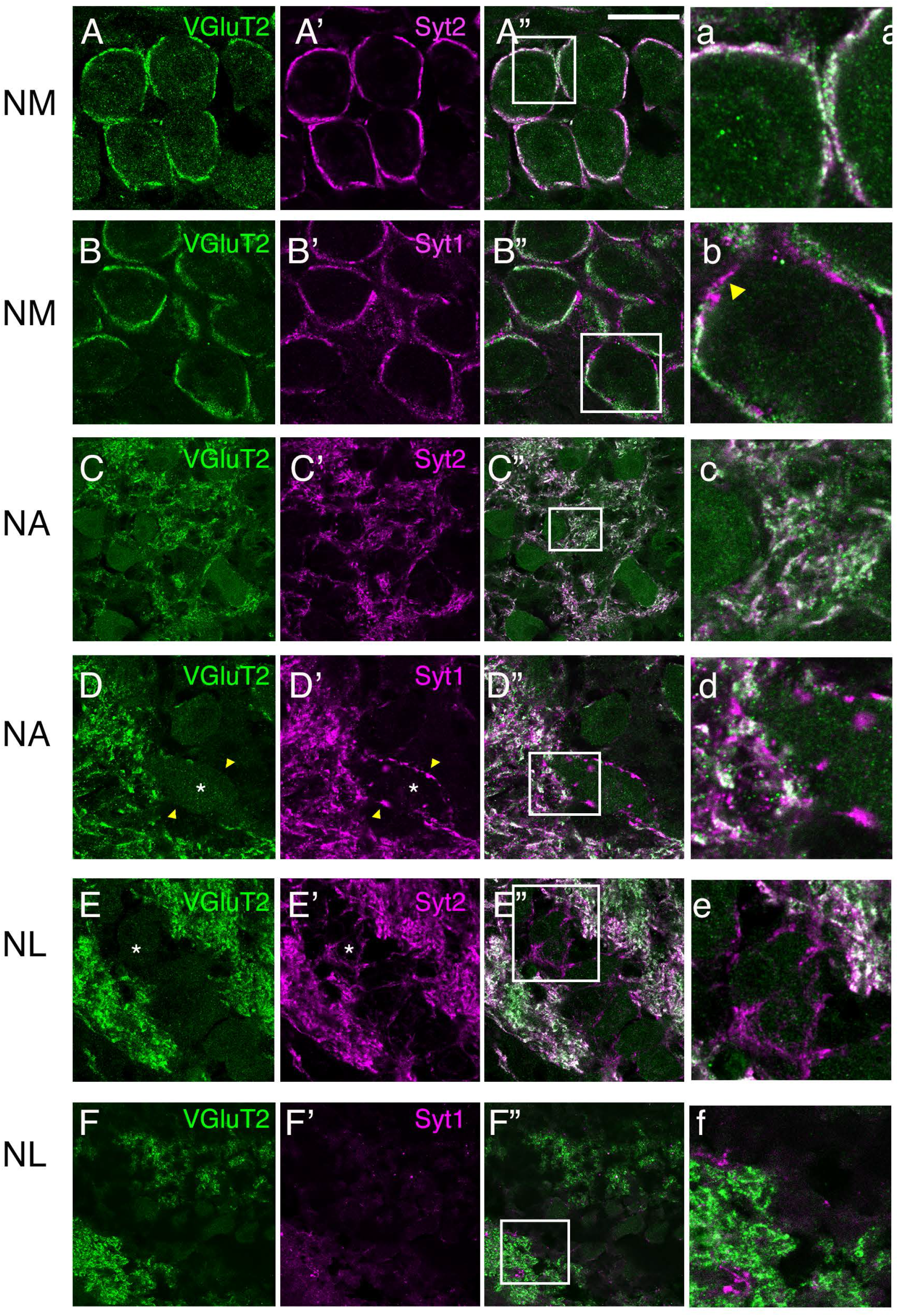
Syt2 and Syt1 were both associated with glutamatergic terminals. Double immunofluorescent labeling of vesicular glutamate transporter 2 (VGluT2, green) with synaptotagmin 1 or 2 (Syt2 or Syt1, magenta, alternating rows), and merged images (right two columns) acquired as confocal single plane images at 63x magnification from NM (A, B), NA, (C,D) and medial region of NL (E,F). Far right panels (a-f) are zoomed in areas (white boxes) from panels A”-F”. Scale bar: 30 µm applies to all panels except a-f. A) VGluT2 and Syt2 co-localized and appeared to be within putative endbulb synaptic endings onto NM cell bodies. B) Syt1 had some areas of non-overlap with VGluT2 (arrowhead). C, D) In NA, VGluT2 densely labeled dendritic neuropil. Syt2 and Syt1 both co-localized with VGluT2 in the dendritic plexus. VGlut2-negative Syt1-positive fibers or en passant synaptic contacts (arrowheads, D-D’) wrapping NA somata (asterisk) were observed. E) VGluT2 and Syt2 both densely labeled the NL dendritic lamina and co-localized there. Syt2-positive, VGlutT2-negative fibers or en passant contacts were observed apposed to cell bodies (asterisk). F) Syt1 expression was extremely sparse in medial NL with little or no co-localization.

The relative lack of anti-Syt1 labeling suggested it was not the primary protein in excitatory terminals in the timing nuclei, but in NM there was some overlap in Syt1 and VGluT2 labeling (Fig. 6B). In NA, anti-Syt1 label and anti-VGluT2 label were mostly overlapping (Fig. 5D). A few Syt1-positive profiles that surrounded cell bodies in NA were VGluT2-negative (arrowheads in Fig. 6D”). In NL, however, anti-Syt1 was so sparse, it too appeared largely unassociated with VGlutT2 (Fig. 6F). In summary, in all three nuclei Syt2 and Syt1 both appear to label excitatory terminals.

### Neither Syt1 nor Syt2 were associated with inhibitory terminals

To determine whether inhibitory terminals also expressed synaptotagmin isoforms, we performed double immunofluorescent histochemistry with antibodies against vesicular GABA transporter (VGAT), a protein responsible for transmitter loading of synaptic vesicles in GABAergic (and glycinergic) terminals. Numerous VGAT+ puncta were observed throughout the auditory nuclei (Fig. 7), similar in pattern to previous avian studies of immunohistochemical labeling of glutamic acid decarboxylase (GAD) (Code et al. 1989; Parameshwaran et al. 2001; MacLeod et al. 2006; Nishino et al. 2008), an enzyme responsible for GABA production. Puncta that were VGAT+ were largely exclusive of either Syt2- or Syt1-label. In NM, the VGAT+ puncta encircled the cell bodies, but double labeling revealed that these puncta did not overlap with and were often exterior to the anti-Syt2 label that resembled the calyx synapse (Fig. 7A). The VGAT+ puncta were also smaller and distinct from the anti-Syt1 label (Fig. 7B). In NA, both anti-VGAT and anti-Syt1 showed the punctate labeling patterns, but the VGAT+ puncta did not appear to colocalize with either Syt1 or Syt2 label (Fig. 7C-D). Similar results were observed in NL: VGAT puncta labeled the dendritic fields as well as ringing the soma in the cell body layer, but did not colocalize with Syt2-positive fibers there (Fig. 7E). Likewise, the Syt1-positive fibers in lateral NL did not overlap with VGAT puncta there (Fig. 7F). These data suggest that neither Syt1 nor Syt2 are the presynaptic calcium sensors for inhibitory terminals in the chick cochlear nuclei or nucleus laminaris.

**Figure 7.**
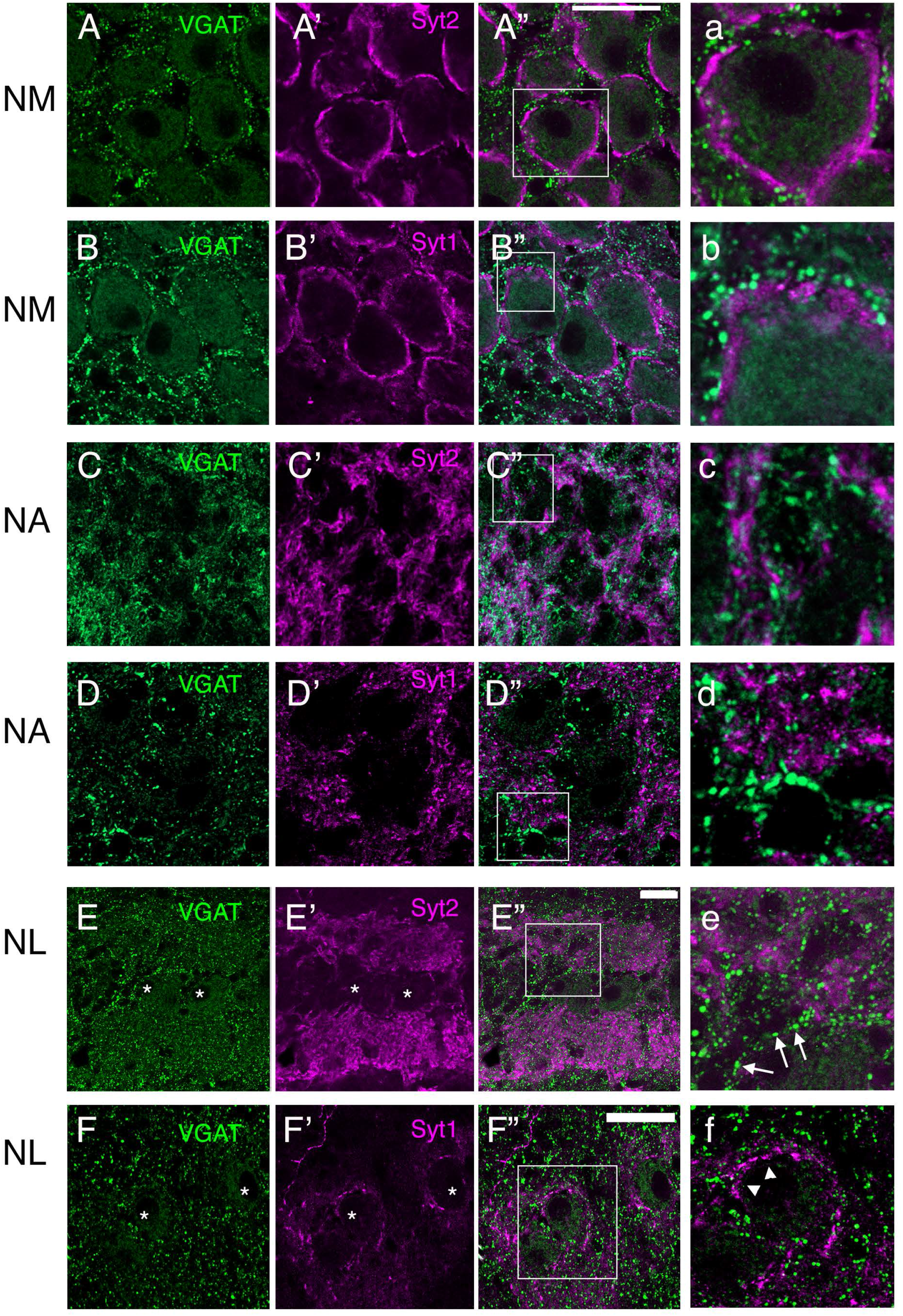
Syt2 and Syt1 were largely not associated with VGAT+ terminals. Panels organized as in Fig. 6 but for VGAT (green) co-labeling with Syt1 and Syt2 (magenta). A) In NM, VGAT positive puncta were distributed in the extracellular matrix between NM cell bodies (A’), and interdigitated with Syt2 positive presumed endbulbs (A’-A”, a), with little to no overlap. Scale bar: 30 µm applies to panels A-D”. B) Syt1 and VGAT also showed almost no overlap in NM. C) In NA, anti-VGAT and anti-Syt2 (and anti-Syt1 in panels in D) both label many puncta throughout NA, but do not overlap. D) In medial NL, VGluT2 and Syt2 both densely labeled dendritic region but were not co-localized. VGAT+/Syt2-puncta lined the somata in the cell body layer (arrows panel e). Scale bar: 30 µm. E) Rare Syt1+ fiber that innervated NL with puncta surrounding the cell body (asterisk) was also VGAT- (arrowheads in panel f).

### Colocalization of Syt1 and Syt2

The labeling patterns for anti-Syt1 and anti-Syt2 antibodies were different from each other, but both appeared to be related to excitatory terminals. To determine whether terminals co-expressed synaptotagmin 1 and 2, we performed double immunofluorescent histochemistry with both the znp-1 (anti-Syt2) and mab-48 (anti-Syt1) antibodies and used isotype-specific secondary antibodies to distinguish between them (see Methods). In NM, many terminals and portions of the calyces that were Syt2-positive were negative for Syt1 (Fig. 8A). In contrast, most Syt1-positive puncta were located within Syt2-positive terminals, or were immediately adjacent to them. In NM this suggests that Syt1 is associated with the endbulbs, but shows subcellular localization, while Syt2 and VGluT2 may be expressed more completely in the presynaptic terminal. In NA, anti-Syt1 and anti-Syt2 did not show much apparent colocalization, but it was difficult to determine whether separate, but adjacent, puncta represented compartmentalized expression or entirely separate terminals.

**Figure 8.**
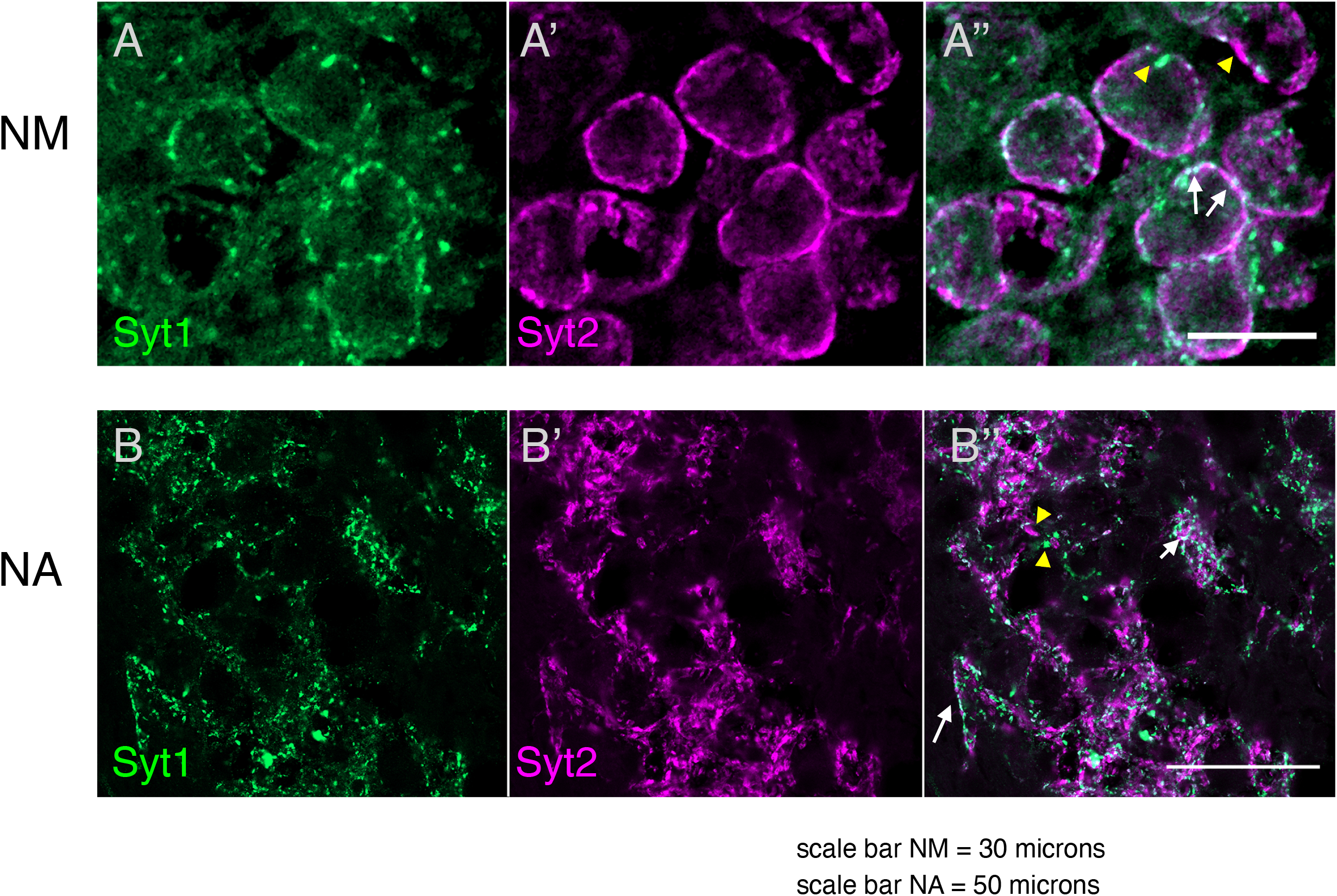
Anti-Syt1 (green) and anti-Syt2 (magenta) label showed substantial but partial overlap in expression in NM (A) and NA (B). In both nuclei, singly labeled (yellow arrowheads) and doubly labeled (white arrows) puncta can be found. Scale bar: 30 µm.

## Discussion

In this study we asked whether different functional pathways in the avian auditory brain stem show differential expression of proteins important for transmitter release. Different isoforms of synaptotagmin have been shown to be differentially expressed in the mammalian and chick cerebellum and retina, as well as mammalian brain stem (Geppert et al. 1994; Fox and Sanes 2007; Sun et al. 2007; Xiao et al. 2010; Kochubey et al. 2011). We investigated the expression of three synaptotagmin isoforms, Syt1, Syt2 and Syt7, in the auditory brain stem nuclei of the chick. Our results demonstrate three main findings: first, that Syt2 was the dominant isoform in all three nuclei, but all three isoforms showed differential expression patterns with the largest differences among nuclei in expression of Syt1. Second, all three isoforms showed mild but statistically significant developmental increases in expression between late embryonic and hatchling ages. Finally, both Syt1 and Syt2 were associated with excitatory synaptic terminals and not inhibitory terminals.

### Differential expression of Syt1, but not Syt2, across auditory brain stem nuclei

The mechanisms responsible for target-dependent regulation of neurotransmission are not fully understood, yet variation in neurotransmitter release dynamics can have a major impact on differential information transmission in circuits (Zucker and Regehr 2002). Previous work in the avian brain stem nuclei, for example, suggests that differences in the short-term plasticity properties of auditory nerve synapses might contribute to the division of auditory information into the “timing” versus “intensity” processing pathways (Cook et al. 2003; Kuba et al. 2005; MacLeod et al. 2007; MacLeod 2011).. Initial release probability is one key property that likely differentiates auditory nerve synaptic transmission onto the two cochlear nuclei, NM versus NA (MacLeod et al. 2007; Ahn and MacLeod 2016) suggesting there could be presynaptic differences in the exocytosis molecular machinery. As is the case with other brain stem areas (Mittelsteadt et al. 2009; Shao et al. 2009; Kochubey et al. 2011) we found that Syt2 is dominant in both avian cochlear nuclei and laminaris, suggesting it is necessary for the rapid, tightly synchronized release. Syt2 is prominent also in the endbulbs of the rodent calyx of Held (Fox and Sanes 2007; Sun et al. 2007; Xiao et al. 2010; Kochubey and Schneggenburger 2011), where Syt1 appears early but then is downregulated (Kochubey et al. 2016). The presence of Syt1 in the caudolateral (low BF) extreme of NL and in NA suggests it could differentially modulate release at these synapses. Alternatively, Syt1 expression could simply represent a redundancy in the molecular machinery for exocytosis.

### Synaptotagmin 7 is sparsely expressed in the auditory brain stem

The presence or absence of short-term synaptic facilitation is a key difference across synapses with different neurotransmitter release dynamics, and the Syt7 isoform has been linked to facilitating synapses (Turecek et al. 2017)(Turecek and Regehr 2018; Huson and Regehr 2020). Thus, we hypothesized that Syt7 may be preferentially expressed at the synapses that facilitated most in the avian brain stem, i.e. those in NA more so than in NM or NL. However, we found that while Syt7 was mostly absent from NM synapses, it was only marginally more expressed in NA and most strongly in NL, contrary to expectation and suggesting an alternative role other than in enhancing synaptic facilitation at these excitatory synapses.

### Synaptotagmin and tonotopy

The avian auditory brain stem nuclei often show gradations in cellular and physiological properties along the tonotopic axis (Rubel and Parks 1975; Jhaveri and Morest 1982; Köppl and Carr 1997; Kuba et al. 2005), some of which can be clearly related to frequency specific computations (Smith and Rubel 1979; Reyes et al. 1996; Agmon-Snir et al. 1998; Kuba et al. 2005; Fukui et al. 2006; Oline and Burger 2014; Oline et al. 2016). Gradients in voltage-gated potassium channels underlie systematic tonotopic changes in electrophysiological properties (Manis and Marx 1991; Rathouz and Trussell 1998; Brew and Forsythe 2005)(Fukui and Ohmori 2004; Kuba et al. 2005). Synaptic release properties also systematically vary with the tonotopic axis, such as synaptic release that is less reliable and depressing in the low BF area of NL (Kuba 2007; Oline and Burger 2014). NA has a tonotopic axis oriented in the dorsoventral orientation (Warchol and Dallos 1990; Köppl 2001; Molea and Rubel 2003), and some expression patterns also show a developmental trajectory that follows this axis (Parks et al. 1997; Kubke and Carr 1998, 2000; Kubke et al. 1999). Our results show that with the exception of Syt1 expression in NL, synaptotagmin expression is not tonotopically varied in the chick auditory brain stem. The stronger labeling in low frequency NL suggest a role for Syt1 in synaptic transmission related to lower temporal precision processing.

### Synaptotagmin expression over late embryonic to early hatchling development

Auditory nerve axons arrive in the chick cochlear nucleus around embryonic day 11 and then undergo morphological and physiological refinement through late embryonic stages and several days after hatching (Jackson and Parks 1982)(see Kubke and Carr 2000 for review). Auditory thresholds drop substantially during late embryonic development prior to hatching and are close to adult levels by P7 (Saunders et al. 1973). Synaptic responses in brain stem neurons also become more reliable and less prone to short-term depression during the same time frame (Brenowitz and Trussell 2001), suggesting alterations to the presynaptic mechanisms of release. Our results show that all three synaptotagmin isoforms demonstrated age-related mild increases in expression levels. Syt2 dominated over Syt1 at all ages, with no evidence at the ages tested for a ‘switch’ from Syt1 to Syt2 as seen in the calyx of Held and other areas (Xiao et al. 2010; Cooper and Gillespie 2011; Kochubey et al. 2016). Strong Syt2 expression was clearly established by E18, although we cannot exclude the possibility that such a switch might occur at earlier embryonic stages. The mild effects of development suggest that the onset of auditory experience only has a minor influence, if any, on synaptotagmin expression.

### Syt1 and Syt2 are associated exclusively with excitatory terminals in the chick cochlear nuclei and laminaris

Substantial differences exist between the presynaptic properties of excitatory and inhibitory neurotransmission in the avian auditory brain stem. Specifically, inhibitory neurotransmission shows greater facilitation and asynchronous release (Monsivais et al. 2000b; Lu and Trussell 2000; Kuo et al. 2012; Burger 2012; Tang and Lu 2012; Curry and Lu 2016; Shi and Lu 2017). Using specific markers of glutamatergic (VGluT2+) and GABA/glycinergic (VGAT+) terminals, we observed an almost complete colocalization of both Syt1 and Syt2 with the glutamatergic but not GABA/glycinergic terminals. The colocalization of Syt1 with VGluT2 and Syt2 further suggests that the punctate nature of Syt1 profiles were not small terminals distinct from the endbulb, but potentially subcellular localization of Syt1 within the calyx. The finding that the inhibitory terminals represented by VGAT labeling were devoid of Syt1 and Syt2 was surprising, given that these isoforms are frequently associated with inhibitory terminals in other systems (Bouhours et al. 2017). The VGAT+ terminals were seen to ring the exterior perimeter of the Syt2 (and Syt1) label that was closely apposed to the cell body, and only sometimes interdigitating with them directly onto the cell body. In chicken NL, VGAT labeling has been shown to be colocalized with the SON projection terminals (Nishino et al. 2008). We found that both Syt2 and VGAT were highly expressed in the dendritic fields, but did not show overlap like Syt2 and VGluT2; in the cell body layer, distinct VGAT puncta outlined the soma (similar to GAD staining) and were clearly Syt2-negative. In NA, Syt2 and VGAT were closely associated, but it was difficult to determine whether these were in anatomically separate profiles or subcellular segregation. The sparseness of Syt1 labeling in NL and its lack of association with VGAT further support the conclusion that Syt1 does not underlie inhibitory neurotransmission. If neither Syt1 or Syt2 underlie inhibitory transmission in the chick auditory brain stem, the next most likely candidate based on its properties and demonstrated ability to support fast synchronous exocytosis would be Syt9 (Xu et al. 2007; Kochubey et al. 2016;). Although Syt9 may be a major component of inhibitory neurotransmission in the mouse striatum (Xu et al 2007), it appears to be largely absent from (or only weakly and transiently expressed in) brain stem (Xiao et al. 2010 ; Mittelsteadt et al. 2009). Determining the identity of the calcium sensor for inhibitory neurotransmission in the chick auditory brain stem will require further investigation.

### Functional role of different synaptotagmin isoforms

Synaptotagmin is an essential protein for presynaptic fast synchronous vesicle fusion and neurotransmitter release. Synaptotagmin 1 and 2 are calcium sensors with similarly low calcium affinity, fast kinetics and appear to be the main isoforms that support rapid exocytosis in association with the SNARE complex. Syt1 and Syt2 have some functional redundancy as both can drive fast synchronous exocytosis, separately or cooperatively (Bouhours 2017)(Xu, Mashimo & Sudhof 2007). Both Syt1 and Syt2 are associated with presynaptic vesicle membrane, facilitate vesicle docking and inhibit membrane fusion until triggered by a large calcium signal, clamp spontaneous release and mediate synaptic vesicle pool replenishment, while Syt1 can help mediate endocytosis as well (Kochubey et al. 2011; Wolfes and Dean 2020). Syt2 has the most rapid release kinetics and vesicle replenishment, which may explain the relative enrichment of Syt2 in hindbrain synapses, and the switch from Syt1 to Syt2 at the calyx of Held (Kochubey et al. 2016). Our results provide support the hypothesis that for highly synchronous release at reliable, temporally precise synapses in the auditory brain stem, Syt2 is required with Syt1 potentially playing a secondary or modulatory role.

Compared to Syt1/2, Syt7 has higher calcium affinity and slower kinetics, may be localized to the cell membrane rather than the vesicle (Adolfsen et al. 2004) and cannot functionally substitute for Syt1/2 in fast synchronous exocytosis (Xu et al. 2007). In addition to being linked to short-term facilitation, Syt7 has also been linked to asynchronous release at some synapses, where it may have a dual function of enhancing synchronous release that is primarily driven by Syt2 (Chen and Jonas 2017). Synaptotagmins are one among many proteins that interact in the presynaptic terminal (Chan and Stanley 2003; Kasai et al. 2012; Jorquera et al. 2012), but expression studies in functionally divergent circuits such as we present here is one step toward linking specific molecular components with function.

## Acknowledgements

Support for this research was provided by a National Institutes of Health grant R01DC10000 to KMM. We also acknowledge Dr. Amy Beaven and the Imaging Core Facility in the department of Cell Biology and Molecular Genetics at the University of Maryland, College Park for assistance with imaging and the use of the Zeiss LSM980 Airyscan 2 supported by National Institutes of Health award 1S10OD025223-01A1.

## Statements and Declarations

The authors have no competing financial interests.

## Materials and Methods

Experiments were carried out on White Leghorn chickens (*Gallus gallus domesticus*) purchased from Charles River Laboratories (Wilmington, MA). All animal procedures were approved by the University of Maryland Institutional Animal Care and Use committee and followed NIH guidelines. Tissue was obtained from chickens of either sex. Animals were perfused at three ages: late embryonic day (E) 17 or 18; hatchling day (day of hatching E21/postnatal day 0) to P1; and week-old P7-8. For simplicity we refer to these age groups as “E18”, “P1” and “P7”. The day of hatch and week-old chickens were deeply anesthetized with isoflurane, while E17/18 embryos were anesthetized by cooling the eggs at -20°C. The chicks were then perfused transcardially with 0.9% saline and heparin, followed by 4% paraformaldehyde (PF) in 0.01 M phosphate buffered saline (PBS, pH 7.4). Brains were postfixed for 18-24 hours, then cryoprotected in 30% sucrose in PBS at 4°C overnight.

### Immunohistochemistry

The primary antibodies were obtained from the Developmental Studies Hybridoma Bank (DSHB; Iowa City, IA), Synaptic Systems (Goettingen, Germany), and Alpha Diagnostic (San Antonio, TX). The primary antibodies used in the present study were (dilutions for enzymatic/fluorescent labeling): mouse anti-synaptotagmin 1 (Syt1) (mab48, DSHB, 1:1k/1:200), mouse anti-synaptotagmin 2(Syt2) (znp-1, DSHB, 1:1k/1:200), rabbit anti-synaptotagmin 7 (Syt7) (Synaptic Systems cat#105173, 1:3k), rabbit anti-VGluT2 (Synaptic Systems, cat#105173, 1:500), and rabbit anti-vesicular GABA transporter (VGAT) (Alpha Diagnostics VGAT11A, 1:500). See Table 1.

The secondary antibodies used were: biotinylated anti-mouse IgG (Invitrogen cat#62-6540, 1:1k), Alexa Fluor 488 donkey anti-rabbit IgG (Invitrogen cat#A21206, 1:200), Alexa Fluor 594 donkey anti-mouse IgG (Invitrogen cat#A21203, 1:200). For Syt1/Syt2 double labeled experiments we used isotype specific, cross-adsorbed secondary antibodies: anti-mouse IgGa, IgGb linked to different fluorophores (Invitrogen Cat#A-21141 and #A-21133, 1:250).

#### Syt antibody specificity

The mab48 antibodies obtained from DSHB originated from immunizing mice with antigen isolated from the cytoplasmic tail of the rat synaptic junction complex protein synaptotagmin-1 (vendor information).The znp-1 antibody was obtained using the full length synaptotagmin-2 full length protein obtained from zebrafish embryos (original reference). The specificity of both antibodies was verified by immunoblot in chickens (Fox and Sanes 2007). Mab-48 and znp-1 label has been extensively characterized in mouse and chick brainstem and cerebellum. The Syt7 antibody obtained from Synaptic Systems was a polyclonal rabbit antibody raised against a recombinant protein corresponding to AA 46 to 133 from rat Synaptotagmin7 (UniProt Id: Q62747). The pattern of label in the chick cerebellum in our study comported with the pattern in mouse (Jackman et al. 2016).

#### VGluT2/VGAT antibody specificity

The VGluT2 (vesicular glutamate transporter 2) antibody (Synaptic Systems 135-403) used in double labeling studies was a polyclonal rabbit antibody raised against a synthetic peptide corresponding to AA 566 to 582 from rat VGLUT2 (UniProt Id:Q9JI12) that is specific for VGluT2 and does not cross-react with VGluT1 (vendor information). VGAT11-A (Alpha Diagnostics) has been previously used in chick auditory brain stem and showed co-labeling with SON descending anterograde traced terminals in NL (Nishino et al. 2008).

### Tissue Processing

Serial sections in the transverse plane were cut on a cryostat at 30-50 µm thickness and collected as alternating sections in triplicate directly onto gel-coated slides. Slides mounted with sections containing the cochlear nucleus angularis, laminaris, and magnocellularis were washed in 0.01M phosphate buffered saline (PBS) then incubated in 10% normal serum in PBS plus 0.5% Triton X-100 for 1 hour. Primary antibody solutions consisted of the appropriate antibody diluted in PBS containing 2.5% normal serum plus 0.5% Triton X-100. Sections were incubated in the primary solution for 1 hour at room temperature then overnight at 4°C. Dilution series were carried out to determine the effective concentrations used in this study. Control sections incubated in the solution where primary antibodies were omitted showed no specific labeling.

Visualization in single antibody experiments was achieved by enzymatic methods using biotinylated secondary antibodies and the avidin-biotin-peroxidase technique. After incubation with secondary antibody solution at room temperature for 1 hour, sections were treated with a hydrogen peroxide solution (1% H_2_0_2_ and 10% methanol in PBS) for 10 minutes to eliminate endogenous peroxidase activity, then incubated in avidin/biotin/peroxidase complex (Vectastain Elite ABC Kit, Vector Laboratories, Burlingame, CA) for 45 minutes. Sections were rinsed several times in PBS, and then the reaction product was visualized with a peroxidase substrate solution (SG Substrate Kit, Vector Laboratories). Slides were washed in PBS and cover slipped with Cytoseal mounting medium (Thomas Scientific, Swedesboro, NJ). Developmental changes in expression were studied using single-antibody IHC experiments were carried out on embryonic and hatchling chickens in independent replicate runs containing animals from three age groups (E18, P1, and P7) processed in parallel and developed identically.

For immunofluorescent labeling, sections were washed and incubated as described above in solutions simultaneously containing two primary antibodies raised in different host species. After rinsing in PBS, sections were then incubated with the solution containing two secondary antibodies tagged with different fluorophores diluted in PBS plus 0.5% Triton X-100 for 2 hours at room temperature. Sections were rinsed, mounted and cover slipped immediately with Prolong Gold containing DAPI mounting medium (Invitrogen Cat# P36935). In experiments using the anti-VGAT primary antibody, tertiary processing was used to enhance the strength of the punctate labeling in which a biotinylated secondary antibody (1:200) was followed by fluorescently conjugated streptavidin (SA-Alexa488, Invitrogen, 1-2 µg/ml).

### Confocal microscopy

Immunofluorescent images were taken on a Leica Sp5X confocal microscope using a 63x oil immersion lens. Some fluorescent images were also taken on Zeiss Airyscan 980 in confocal mode. Images were composited and adjusted for brightness and contrast in ImageJ.

### Optical Density Analysis

Optical density analysis was performed on sections processed for single antibody immunohistochemistry against synaptotagmin 1, 2 and 7 and visualized by enzymatic methods using peroxidase precipitation. Images were taken using the Neurolucida software (Microbrightfield) on an Olympic upright microscope using brightfield and 10x or 20x objective for each section, camera gain and exposure were adjusted to maximize the range of values between 0 and 255, and then using the default settings for that range. Several fields of view were collected to create a montage of a brain stem dorsal quadrant with identical image acquisition settings. Images were imported into ImageJ (FIJI), regions of interest (ROI) outlined and mean grayscale values measured within each ROI. Optical density measurements were made of whole nucleus regions in each section unless otherwise noted. Nuclear borders were determined by the labeling itself and visible contours of the nuclei in brightfield surrounded by fiber tracts, but were also consistent with comparable sections labeled with calretinin or SV2. Medial, middle, and lateral (or dorsal and ventral) regions were determined by dividing the nucleus area in to thirds across the mediolateral (NM and NL) or dorsoventral (NA) axis. In NA where there was a clear dorsolateral region, it was included in the general dorsal ROI. A more detailed analysis of NL used the anatomical mapping method that more accurately follows the rostromedial to caudolateral orientation of the frequency mapping, described by Kuba et al. (2005). Briefly, four sequential transverse sections containing NL were divided into 10 sectors, labeled 2-11 (sector 1, at the rostral pole was omitted): rostralmost section 1 contained sectors (from medial to lateral within section) 2 and 3, section 2 contained sectors 4-6, section 3 contained sectors 7-9, and caudalmost section 4 contained sectors 10-11.

### Statistics

All data are presented as mean ± standard error unless otherwise noted. Statistical analyses of OD measurements were carried out using GraphPad Prism (v.9.0.0). Statistical comparisons of tonotopic gradient differences within nuclei were made using non-parametric tests and within section paired or matched data.

